# Imaging and mechanism of DNA-DNA recognition mediated by divalent ions

**DOI:** 10.1101/2025.09.11.675610

**Authors:** Thomas E. Catley, Victor Velasco-Berrelleza, Daniel E. Rollins, Alice L. B. Pyne, Agnes Noy

**Author notes:** Authors contributed equally (Thomas E. Catley, Victor Velasco-Berrelleza).

## Abstract

In the cell, DNA must be tightly packed to facilitate its organisation into the nucleus, where recognition of homologous sequences underpins key processes such as recombination. Yet the structural basis of DNA–DNA pairing remains unknown. Here we combine high-resolution AFM and atomistic MD simulations to provide the first direct visualisation of DNA pairing in the presence of divalent ions. We show that strongly paired DNAs often achieve groove-to-groove alignment, driven by ionic bridges connecting the minor grooves of the two duplexes. These contacts are further stabilized by sequence-specific interactions, whose strength and specificity vary with the ion type. This mechanism of ion-mediated groove alignment provides a molecular framework for the long-hypothesized “helical alignment” model, in which homologous recognition is facilitated by preserving structural register between the two helices. Together, our findings reveal a fundamental principle by which divalent ions promote DNA–DNA pairing, with broad implications for chromosomal condensation and genome organization.

## Introduction

The condensation of DNA is essential for genome compaction ^1–4^, as well as for DNA nanotechnological applications, such as DNA origami^5^. Furthermore, the association of homologous double stranded DNA (dsDNA) molecules has been observed in early meiosis prior to the action of specialized recombination machinery^6–8^. Homologous chromosome pairing without recombination has also been detected in *Drosophila melanogaster*^9,10^ and in mammals, which has been associated with processes such as X-chromosome inactivation^11^ and cancer^12^.

The molecular mechanisms that underlie the recognition of homologous sequences are still unknown. It has been investigated using the Repeat-Induced Point Mutation (RIP) system in *Neurospora crassa*, which provides the most compelling evidence of this effect *in vivo*^13^. In this organism, experiments have revealed that homologous trinucleotides are sufficient for pairing when they are repeated every 11 bp^14^. These findings indicate that homology recognition could be facilitated by the direct interaction between DNA duplexes, as this frequency is in accordance with the helicoidal periodicity of DNA. The mediation of RNA and proteins would secure this pairing, although an initial recognition would still be necessary^15,16^.

It is well known that dsDNA molecules can condense in the presence of polyamines such as spermidine^3+^ and spermine^4+ 17^. These polycations are present at a millimolar concentration in most eukaryotic cells and are known to participate in chromatin condensation^18^. Molecular dynamics simulations revealed that this condensation is mediated by the bridging of polyamines between DNA molecules and is greatly influenced by sequence (being stronger in AT-rich sequences than GC-rich)^19,20^. Although this model could explain homologous sequence recognition via the alignment of sticky sequence patches, it cannot account for the dependence of partial homology on 11-bp periodicity^14^, due to polyamine bridging occurring regardless of the orientation of the two duplexes.

A variety of experimental techniques have provided evidence that divalent ions at physiological concentrations can also induce attraction between DNA duplexes, particularly when homologous sequences are coupled^21–26^. While monovalent salts seem also to promote pairing, this effect would generally require significantly longer DNA molecules (>1 kbp) compared to the shorter fragments (typically <100 bp) sufficient for divalent ion-mediated attraction^23,24^. A proposed model for interpreting these observations has been the “electrostatic DNA zipper”, or helical coherence model, originally introduced over two decades ago and widely explored since^27–29^. In this framework, counterions bind or condense to DNA grooves, enabling electrostatic interlocking between the positively charged grooves of one duplex and the negatively charged backbone of another^27^. Homologous sequences would then stabilize this interaction by preserving sequence-dependent variations in helical pitch, thus maintaining groove–strand registry along the molecules^29^. Alternative hypotheses have suggested that direct inter-duplex hydrogen bonding across DNA grooves may contribute to recognition, potentially through transient noncanonical conformations such as C-DNA– like structures^30^. However, such conformations have not been experimentally observed in solution^31^. Despite all these efforts, the precise molecular mechanism of DNA–DNA pairing remains elusive and lacks direct experimental visualization.

In this study, we present the first direct imaging of DNA pairing in a solution of divalent ions using high-resolution Atomic Force Microscopy (AFM). We observe that DNA grooves are strongly aligned across the duplexes, and that the same register is maintained consistently throughout the pairing in line with the zipper model. These results are supported by molecular dynamics simulations that reproduce spontaneous DNA-DNA self-assembly when solvated by divalent cations. Our simulations show that DNA-DNA contacts are stabilised by an ion-bridging mechanism which is facilitated by specific coordination sites within the minor groove. Both AFM and MD data reveal that these coordination sites occur at non-random positions, indicating that the pairing mechanism is intrinsically sequence-dependent and varies according to the specific divalent ion. This sequence-specific localization is consistent with previous observations in *Neurospora crassa*, where DNA motifs were found to recur periodically in phase with the helical pitch^14^. We propose that these sites act as anchoring points for the initiation and propagation of pairing, a process likely facilitated by sequence homology. Taken together, our results provide a plausible explanation for the mechanism of homologous recognition.

## Materials and Methods

### Sample Preparation for AFM

DNA samples were deposited on a freshly cleaved muscovite mica surface following a published protocol^32^. For circular DNA, 17 ng of DNA was deposited into 20 µL of immobilisation buffer (10 mM NiCl_2_, 20 mM HEPES, pH 7.4). For linearised DNA, 3-8 ng of 339 bp linearised DNA was deposited into 30 µL of immobilisation buffer. The immobilisation buffer consisted of either 10 mM NiCl_2_, 10 mM CaCl_2_, or 10 mM MgCl_2_, along with 20 mM HEPES. The pH of all buffers was adjusted to pH 7.4 with NaOH, resulting in a final Na^+^ concentration of 8.5 mM. This was incubated for 30 minutes before being washed 4 times in 30 µL of the same buffer. After the final wash, a final 30 µL of 10 mM NiCl_2_ was added to the surface to ensure stable imaging.

All AFM measurements were performed in liquid following a published protocol^32^. Imaging was carried out in PeakForce Tapping mode either on a Multimode 8 AFM system (Bruker) or a Dimension XR FastScan (Bruker). For imaging on the Multimode 8, PeakForce HiResB cantilevers (Bruker) were used and for imaging on the Dimension FastScan, FastScan D cantilevers (Bruker) were used. The PeakForce Tapping amplitude was set to 10 nm, the PeakForce Tapping frequency to 4 KHz on the Multimode 8 and 8 KHz on the Dimension FastScan. The PeakForce setpoint was maintained in the range 10-20 mV, corresponding to peak forces of <70 pN. The number of samples per line was selected to maintain a resolution of at least 1 px/nm.

### AFM image processing and analysis

All AFM images were processed using TopoStats, a Python pipeline for the automated processing and analysis of AFM data (https://github.com/AFM-SPM/TopoStats)^33^. TopoStats can be configured using a configuration file. In this instance, the default settings were used with 3 changes. Firstly, the upper threshold for the filtering and grain detection stages was changed depending on the sample type to obtain the highest quality flattening, and secondly the height range for the images was set to [-3, 4] in order to produce images where height differences were more easily visible. The absolute area thresholds were also adjusted to filter noise and anomalies of varied size in the data. The configuration file can be found with the dataset. Processing was carried out using the following steps: an initial flatten removes image tilt by subtracting a plane, then, to remove line-to-line variation from the raster scan process, the median of each row is subtracted. Finally, to remove bowing effects, a horizontal quadratic polynomial is subtracted. Finally, a gaussian blur of sigma = 0.5 pixels was applied to reduce high frequency noise in the image.

To quantify the DNA recognition effect, the image analysis software FIJI^34^ was used to import the processed image files and manually trace the DNA strands. The coordinates of the strands were then passed to a custom python script which determined whether condensation was taking place by measuring if the central axes of the strands passed within 6 nm of one another. This value was chosen to account for the tip broadening effects commonly seen with AFM imaging. The classification of the alignment of the condensed DNA strands was then carried out by visual inspection.

### MD simulations

Four 28 bp-long segments were extracted from the DNA 339 bp fragment (see supplementary information for sequence and positions). Two C:G bp were added to each end in order to prevent melting during simulations. The DNA duplexes, 32 bp in total length, were constructed using the NAB module in Amber20^35^. Two DNA duplexes were placed in parallel separated by 3.5 nm according to the central DNA axis and orientated in two different azimuthal angles, 0 and 180 degrees (see Fig. S1). DNAs were represented by the parmbsc1 forcefield^36^, solvated by TIP3P octahedral boxes with a buffer of 1.5 nm.

Initial simulations were performed using 100 mM concentrations of Ni(Cl)_2_, Mg(Cl)_2_, Ca(Cl)_2_ and 200 mM of KCl, with ions randomly distributed throughout the solvation box. Subsequent simulations were conducted under physiological conditions (150 mM KCl and 20 mM Mg(Cl)_2_ or Ca(Cl)_2_^37^ using two distinct protocols: (i) a fully random distribution of all ions; and (ii) placement of Mg2+ or Ca2+ in the most energetically favourable positions while KCl pairs randomly distributed. In a final set of simulations, the negative charge of the DNA was fully neutralized by Mg^2+^ or Ca^2+^ placed in the most energetically favourable positions, with additional 150 mM KCl randomly distributed. In all simulations, the final ionic strength was adjusted using the Split method^38^.

Ni^2+^ was simulated with dummy model parameters, which have demonstrated the ability to replicate experimental solvation free energies, ion–oxygen distances, and coordination numbers^39^. The dummy ion model was initially developed for transition metal ions such as Ni^2+^ to resolve the challenge of precisely correlating solvation energies and ion-oxygen distances using a simple Lennard-Jones sphere with a central charge^40^. This type of ions are characterized for presenting higher hydration energies compared to an alkaline-earth ion of equivalent size. The model redistributes the charge from the centre by incorporating extra dummy atoms to properly represent the electronic configuration surrounding the ion. Mg2^+^ and Ca2^+^, and the associated Cl^−^, were modelled using the 12-6-4 Lennard-Jones model from Li and Mertz^41^, recognized as among the most accurate parameters for describing alkaline-earth ions^42^. The monovalent ions K+ and Cl-were represented by the widely used Dang and Smith parameters^43^.

Following minimization and thermalization (T = 298 K), a 300 ns equilibration step was conducted with positional restraints of 100 Kcal/mol on DNA atoms to allow the ions to equilibrate around the DNA molecules. This equilibration step was extended to 500 ns for the simulations under physiological solvent conditions where ions were distributed randomly to provide extra time for ion equilibration. The final structures were subject to 200 ns of productive restriction-free MD simulation at constant temperature (298 K) and pressure (1 atm) using periodic boundary conditions, particle mesh Ewald and an integration time step of 2 fs, following our standard protocols^44^. The last 100 ns were used for the subsequent analysis with the objective to characterize self-aggregation. Table S1 provides an overview of all simulations, totalling 40 and covering a cumulative time over 20 µs.

### Simulation analysis

To extract representative structures from all simulations, we applied the hierarchical agglomerative clustering based on the average linkage algorithm as it is implemented in cpptraj^45^. The root-mean-squared deviation (RMSd) between all frames from the 200-ns restriction-free simulation was used as a distance metric. The frame closest to the centroid of the most populated cluster from the last 100 ns was selected as the representative structure. The number of clusters was chosen so that each had a clearly distinct arrangement between the two duplexes. They ranged from 2 in the most condensed pairings to 6 in the least condensed ones, illustrating the process of pairing (Fig. S2 and S3).

To measure the intermolecular distance between two DNAs, we used our own software, WrLINE^46^, which extracts the central axis of each DNA duplex with one point per each bp. Next, we generated a matrix that contained all distances between axes points across duplexes and identified the diagonal with the smallest trace (Fig. S2 and S3). This was done to determine the optimal alignment between the two molecules. The average intermolecular distance was calculated by considering only the 21 lowest distances, corresponding to two DNA turns, to exclude uncondensed ends. To quantify alignment and distance between minor grooves across duplexes, a contour along each duplex’s minor groove was constructed using the center points between phosphate atoms. Lines were fitted to each groove contour considering a sliding window of three points. Angles and distances between lines of the two duplexes were calculated. As previously done, matrices comprising all angles and distances were created to determine the lowest trace. The average angle and distance were calculated using only the two lowest values, which corresponded to two DNA turns, following the same criteria as before. The deviation from planarity was calculated through the minimal perpendicular distances between the WrLINE molecular contours and best-fitting planes for each individual frame of simulations using our software SerraLINE^47^. Ion densities around DNA duplexes were determined by the program Canion^48^. We identified ions that form DNA-ion-DNA bridges based on their distance within the second solvation shell of the two duplexes, with a cutoff of 5 Å.

## Results

### High-Resolution AFM reveals alignment of major and minor grooves in paired dsDNA

High-resolution images of DNA condensation were captured while imaging a sample containing DNA 250-350 bp minicircles and linear fragments in the presence of 10 mM NiCl_2_. The use of Ni^2+^ as counterions allows for optimal spatial resolution by AFM, allowing the detection of sub-molecular details necessary to resolve DNA grooves^49,50^. These images revealed the close packing of DNA duplexes where the major and minor grooves are highly aligned (Fig. 1). We observed several instances of this alignment (Fig. 1a), whereby the DNA backbones appear to be in contact, and the major (pink star) and minor (green triangle) grooves lie alongside each other. This is exemplified in Fig. 1b, which shows near-perfect alignment of the grooves between the two paired duplexes. We verify this alignment by measuring height profiles along each DNA backbone (Fig. 1c), finding that the reduction in height corresponding to the major and minor grooves^49^ occur in the same position along each duplex. These images are consistent with the previously proposed electrostatic zipper model whereby the action of divalent ions align the grooves on adjacent duplexes^27^, providing the first direct visualisation of this effect.

**Figure 1.**
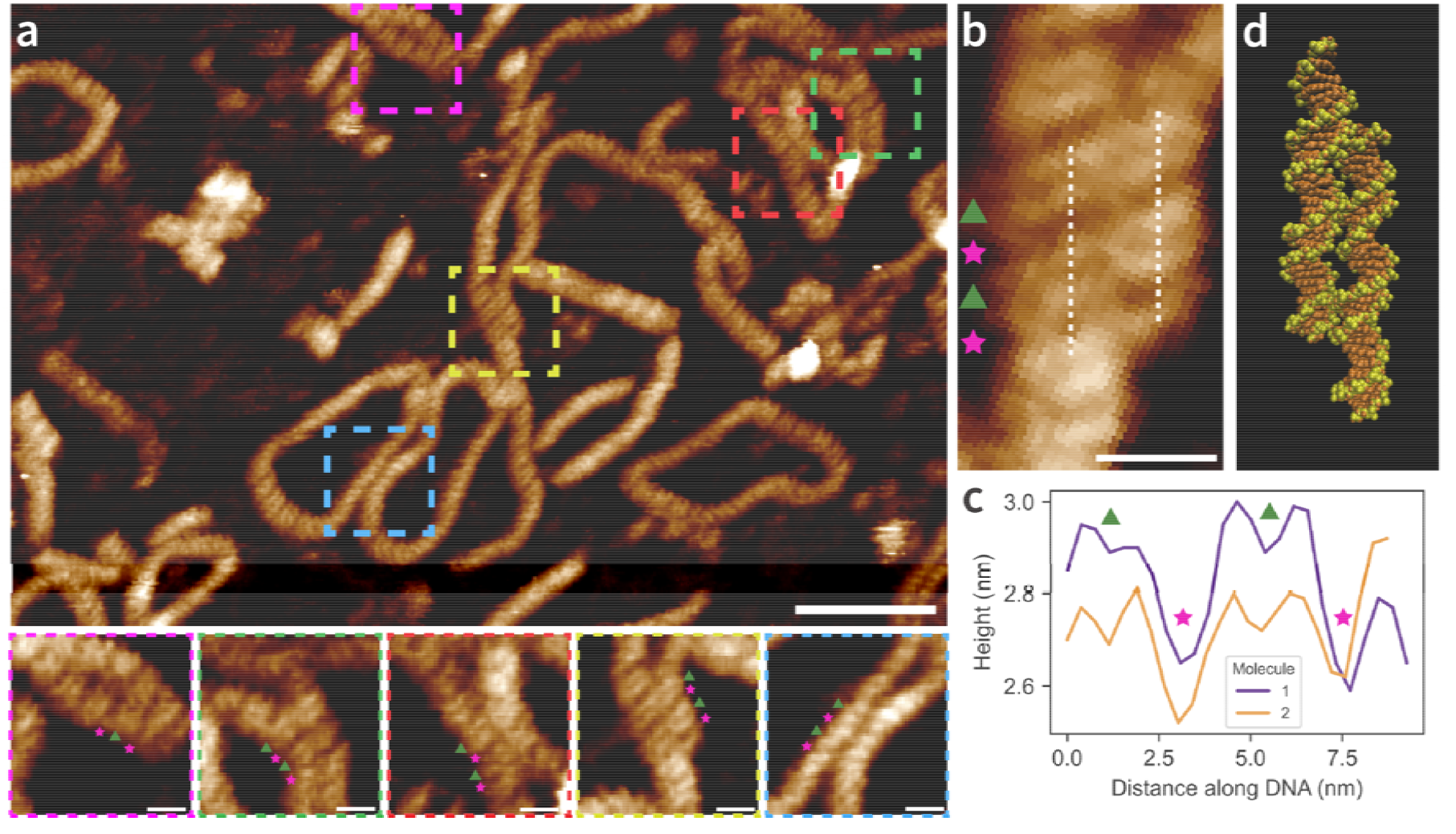
DNA-DNA recognition aligns major and minor grooves between duplexes. **a)** High resolution AFM images of 251 bp DNA showing several examples of strong alignment of the major and minor grooves between duplexes within close proximity. Scale bars = 30 nm (large image) and 5 nm (small images). Height scales = −0.5 to 2 nm (large image) and 0 to 2 nm (small images). Green triangles indicate location of the minor grooves and pink stars indicate the location of the major groove. **b-c)** Parallel height profiles taken along the adjacent duplexes highlight the alignment of the major and minor grooves. **d)** MD simulations of linear fragments ~30 bp long present spontaneous DNA-DNA pairing, exhibiting similar alignment in the grooves.

### DNA pairing is observed in different configurations and locations along the duplex depending on ionic conditions

To further investigate how pairing occurs, we linearised a 339 bp minicircle with the restriction enzyme NdeI. The resulting linear molecules allowed us to determine whether pairing formed at specific positions along the DNA. Given that resolution of the major and minor grooves of the DNA is not always achievable, large area AFM scans were taken in the presence of either 10 mM NiCl_2_, 10 mM CaCl_2_ or 10 mM MgCl_2_ (Fig. S4) to quantify the pairing effect in different ionic conditions over a large dataset (NiCl_2_ = 56 paired out of 809 total molecules, CaCl_2_: 54/906, MgCl_2_: 52/925).

We classified interactions as “pairing” if the distance between the centre of the DNA backbones was ≤6 nm, since AFM tip convolution resulted in an average measured DNA width of 6 nm (Fig. S5). These images revealed that the DNA duplexes interact in multiple configurations, which we categorised and subsequently analysed using semi-automated tracing (see methods) to quantify both the length and location of pairing (Fig. 2). The “mid-symmetric” and “mid-asymmetric” states were characterized by the close contact of the central 60% of the molecule, resulting in symmetric or asymmetric tails between the two duplexes. The “end-to-end” category includes molecules with alignment of terminal 20% of two DNA duplexes, while the “mid-to-end” category includes pairs where a fragment’s terminus interacts with the central portion of the second duplex (Fig. 2a). Although the direction of pairing cannot be determined from the AFM images, we can establish that fully homologous pairing is only possible in the two “symmetric” states (i.e., “mid-symmetric” and “end-to-end”).

**Figure 2.**
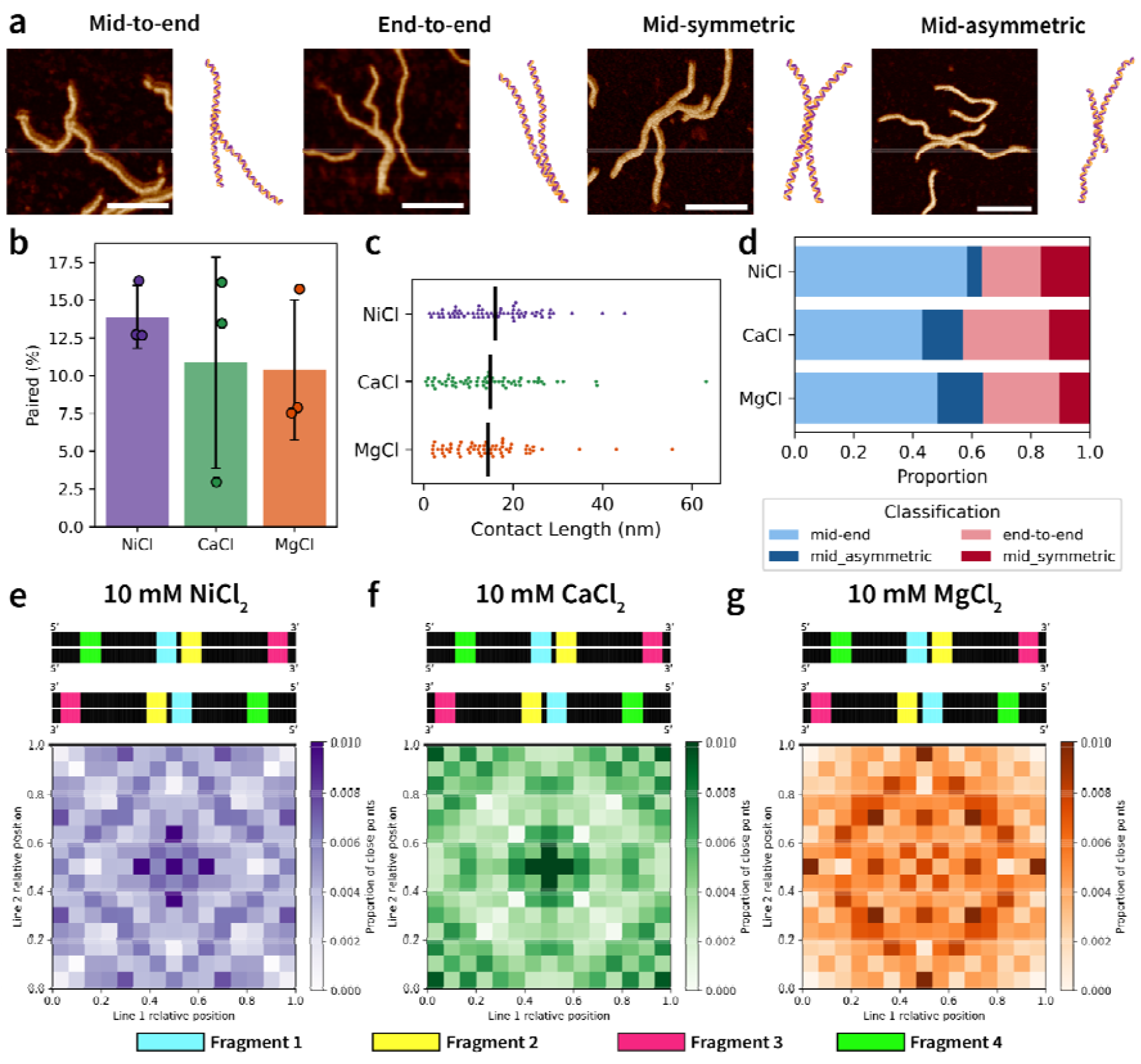
DNA duplexes show preferential pairing configurations. **a)** Classification of pairing conformation types observed in AFM images. Scale bar = 50 nm, Height scale = −1 to 3 nm. **b)** Proportion of total molecules found in a pairing event across different ionic conditions (N = 3 independent replicates). **c)** Total contact length between DNA duplexes during pairing events. **d)** Distribution of pairing classifications for 339 bp fragments immobilized in each ion condition. 2D pairing maps along the DNA molecule in 10 mM NiCl_2_ (**e**), 10 mM CaCl_2_ (**f**), and 10 mM MgCl_2_ (**g**), where darker regions indicate higher frequencies of close pairing. Values are mirrored around the 0.5 midpoint, as duplex directionality cannot be determined from AFM measurements. The schematics above panels (**e-g**) illustrate the positions of the fragments used in MD simulations relative to the 339 bp linear DNA sequence, accounting for both possible orientations. Statistical significance was assessed using a Kruskal-Wallis test with Dunn’s pairwise comparison in (**b**) and (**c**), revealing no significant differences between groups. Differences between the distributions in (**d)** and the heatmaps in (**e–g**) were evaluated using pairwise Pearson’s chi-squared tests for independence.

We found that the mean percentage of molecules imaged in a paired state was not significantly different between the different ions (NiCl_2_: 14 ± 2%, CaCl_2_: 11 ± 7%, MgCl_2_: 10 ± 5%) (Fig. 2b). Additionally, there was no significant difference between the length over which the DNA duplexes remained in contact during a pairing event in the different conditions (NiCl_2_: 16 ± 9 nm, CaCl_2_: 15 ± 11 nm, MgCl_2_: 14 ± 10 nm) (Fig. 2c). This indicates that the ionic conditions tested did not significantly influence the strength or extent of duplex pairing. Furthermore, the data are consistent with the high-resolution AFM measurements (Fig. 1a) showing duplexes remaining in contact over 3– 4 helical turns (~11–15 nm).

When evaluating the proportions of pairing states, we found that the non-homologous conformations (i.e. the “mid-to-end” and “mid-assymetric”) were the most prevalent in all ionic conditions, accounting for 63% of the pairings in NiCl_2_, 57% in CaCl_2_, and 64% in MgCl_2_. (Fig. 2d). As we would expect dominant configurations to arise from homologous pairing (i.e. “end-to-end” and “mid-symmetric”), these results suggest that there are other factors that enable close pairing of the duplexes in these conditions. These could include a distinct propensity of the central and terminal regions in the DNA to meet, or ion mediated matching driven by the presence of particular sequence motifs.

Our semi-automated tracing pipeline allows to detect the position of pairing events along both duplexes, which provides richer information than simple classification of configurations. From these, an average “contact map” can be constructed which highlights the regions along the duplex that come into close contact most frequently (Fig. 2e-g). In 10 mM NiCl2, this contact analysis shows a pairing hotspot at the centre of the duplex (Fig. 2e), between 30 and 70% from the end, where there is an elevated degree of pairing. This indicates the possibility of sequence motifs that are driving pairing in this particular region. As with the NiCl2, the pairing for CaCl2 (Fig. 2f) is concentrated at the centre of the duplex, indicating that the pairing is also strongest in this region. However, in MgCl2 (Fig. 2g) the pairing is observed to be more evenly distributed along the duplex. The pairing heatmaps revealed statistically significant differences between all ionic conditions (Pearson’s chi-squared comparisons: NiCl_2_ vs. MgCl_2_, χ^2^ = 3300; NiCl_2_ vs. CaCl_2_, χ^2^ = 6300; CaCl_2_ vs. MgCl_2_, χ^2^ = 8800), indicating that the spatial distribution of pairing events along the duplex varies depending on the ionic environment. Because electrostatic repulsion is arguably higher when the middle sections of two DNA duplexes come together than when the ends meet^51^, we hypothesize that certain sequence motifs near the center of the DNA segment may aid in pairing in the presence of NiCl_2_ and CaCl_2_, but to a lesser extent in MgCl_2_.

### Atomic MD simulations reveal DNA-DNA pairing consistent with AFM observations

To describe the mechanism of DNA-DNA condensation observed by high-resolution AFM images, allatom MD simulations were performed with Ni2+ as the divalent counterion. Four ~30 bp-long fragments were selected from the experimental sequence to represent areas which showed different propensity for pairing by AFM (Fig. 2). We chose two fragments from the middle to capture the best pairing region (fragments 1 and 2) and two at the ends to examine sequences with less propensity to pair (fragments 3 and 4). We simulated the four homologous pairs by combining each sequence with itself. In addition, we simulated four non-homologous pairs by mixing sequences from the central region (fragments 1 and 2) and across regions (fragments 1 with 4, 2 with 3 and 2 with 4).

To observe whether all-atom MD simulations could reproduce spontaneous self-assembly, MD simulations were initiated with two dsDNA fragments positioned 3.5 nm apart in a parallel configuration. A high concentration of Ni^2+^ (100 mM) was employed to facilitate a sufficient ion population between the duplexes, which is otherwise limited by slow exchange kinetics with DNA. Two independent simulations were run for each pair, with the relative orientations of the duplexes altered by 180 degrees (see Fig. S1). We fixed the coordinates of the DNAs for the initial 300 ns to allow the ions to equilibrate around them and then set the DNAs free for 200 ns.

The pairs of dsDNA fragments interacted with each other in almost all simulations containing Ni^2+^, even if they were not homologous (Fig. 3 and S6). This is consistent with the observation of anti-parallel pairings by AFM, where homology cannot be the only driver. The average distance between the molecular axes of the two duplexes fell below 2.5 nm in most of the Ni^2+^ simulations, approaching the duplex’s width (~2 nm) and indicating highly closed condensation (Fig. 3a, b). Structural characterisation showed that contact spots between duplexes occur when the two minor grooves are oriented towards one another, facilitated by high local densities of Ni^2+^ ions (Fig. 3c). We measured distances and angles between neighboring minor grooves across duplexes and noted a correlation of 0.81 and 0.40, respectively, with the intermolecular distance. These correlation coefficients were computed excluding replica 2 of fragments 1-4, which exhibits minimal condensation (see Fig. S6). This indicates that the most closely paired DNAs tend to be those with least separation between minor grooves on neighbouring molecules and with high groove alignment. Structures presenting these characteristics resembled those captured by high-resolution AFM (Fig. 1b, d and Fig. 3c).

**Figure 3.**
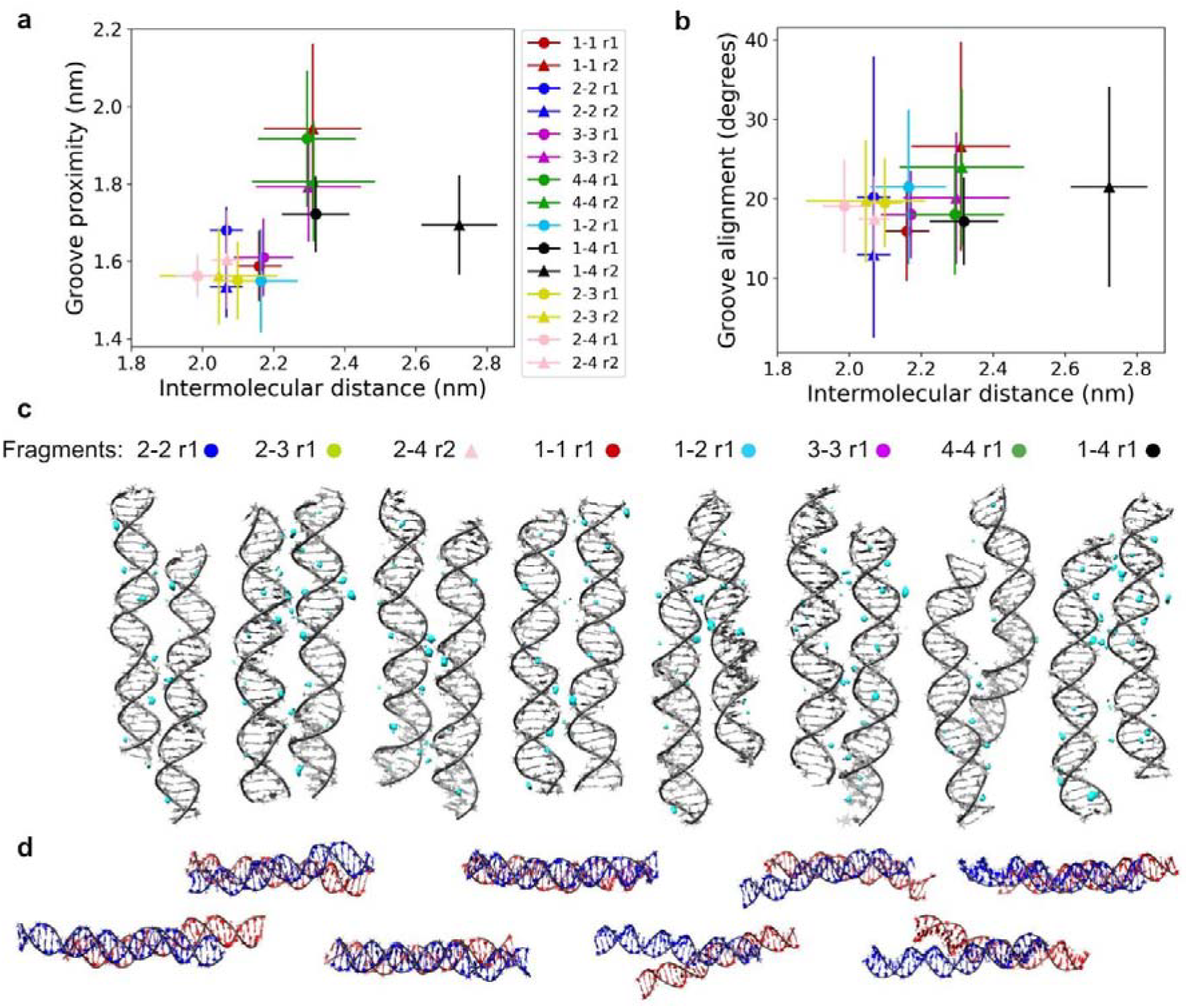
MD simulations reveal spontaneous DNA-DNA pairing mediated by Ni^2+^ resembling high-resolution AFM images. **a-b**. Strength of intermolecular DNA pairing determined by the distance (**a**) and angle (**b**) between minor grooves of adjacent duplexes. DNA pairings are coloured by the fragments that constitute them (1, 2, 3 or 4) with replicas 1 and 2 (r1 and r2) differentiated by shape. Replica 2 for pair 1-2 did not present condensation and so is not included. Values reported are averages over all frames from the last 100 ns of each simulation with the corresponding standard deviations (error bars). **c**. Representative structures for each DNA pair, ordered from lowest intermolecular distance (left) to highest (right). Ni^2+^ density maps an occupancy of 25 times or greater the bulk concentration (in cyan). The replica with the smallest average distance between duplexes is shown, and if they are similar, the most planar one (Fig. S8). The remaining replicas are presented in Fig. S6. **d**. Representations of planarity corresponding to the paired DNAs above with the front duplex in blue and the rear duplex in red. The juxtaposition of DNA double helices can be predominantly planar (Fig. S6 and S8), which explains the good agreement between AFM on a surface, and by MD simulations in solution. When the pairing adopts a more three-dimensional shape the resultant crossovers can exhibit either left-handedness or right-handedness, often involving the molecular ends, which could potentially be caused by the limited length of our DNAs (Fig. S6). Consequently, we concentrated on the most planar pairs to examine in detail the mechanism through which divalent ions could facilitate DNA-DNA condensation.

We performed multiple replicates across various sequences and initial orientations in 200 mM KCl without observing DNA–DNA pairing in any instance (Fig. S7). These results are consistent with previous experimental findings indicating that monovalent salts fail to induce pairing in relatively short DNA molecules (<500 bp)^21,22^, with condensation only occurring for significantly longer fragments (>1 kbp)^23,24^.

### Sequence motifs drive DNA-DNA pairings

We observed that the most densely packed DNAs involved fragment 2 from the central region, with the pairings being either homologous or non-homologous (Fig. 3), consistent with the AFM images shown in Fig. 2. The ion density maps surrounding the simulated DNA structures revealed a series of high-density spots between the two duplexes, corresponding to well-positioned ions (Fig. 3c). Some of these ions were simultaneously interacting with both duplexes, thereby producing a DNA-ion-DNA bridge (Movies S1-S4). We identified the ions that produced the most stable bridges (i.e. the ones maintaining the interaction for the longest time in our simulations) and observed that they were predominantly positioned inside the minor groove of the GTAC sequence motif (see Table S2). A closer look at the simulations showed that nickel ions contact the partially negatively charged N3/O2 atoms of the central A/T bases. These are held in place by the large NH3 group from the nearby CG base pair, creating a binding pocket (Fig. 4), supported by additional interactions with the backbone of the other DNA duplex. This mechanism explains the high stability of these bridging ions, which had residence times exceeding 60 ns (Table S2). The two facing minor grooves formed a pairing site that was stabilized by two bridging ions, one in each minor groove, and, on some occasions, by a third ion placed between the backbones of the different duplexes, which was considerably more dynamic than the other two (Fig. 4 and Table S2). We observed that the GTAC motif participates in all the main pairing sites, with the other sequences involved containing stable ions located opposite to it (Fig. S9).

**Figure 4.**
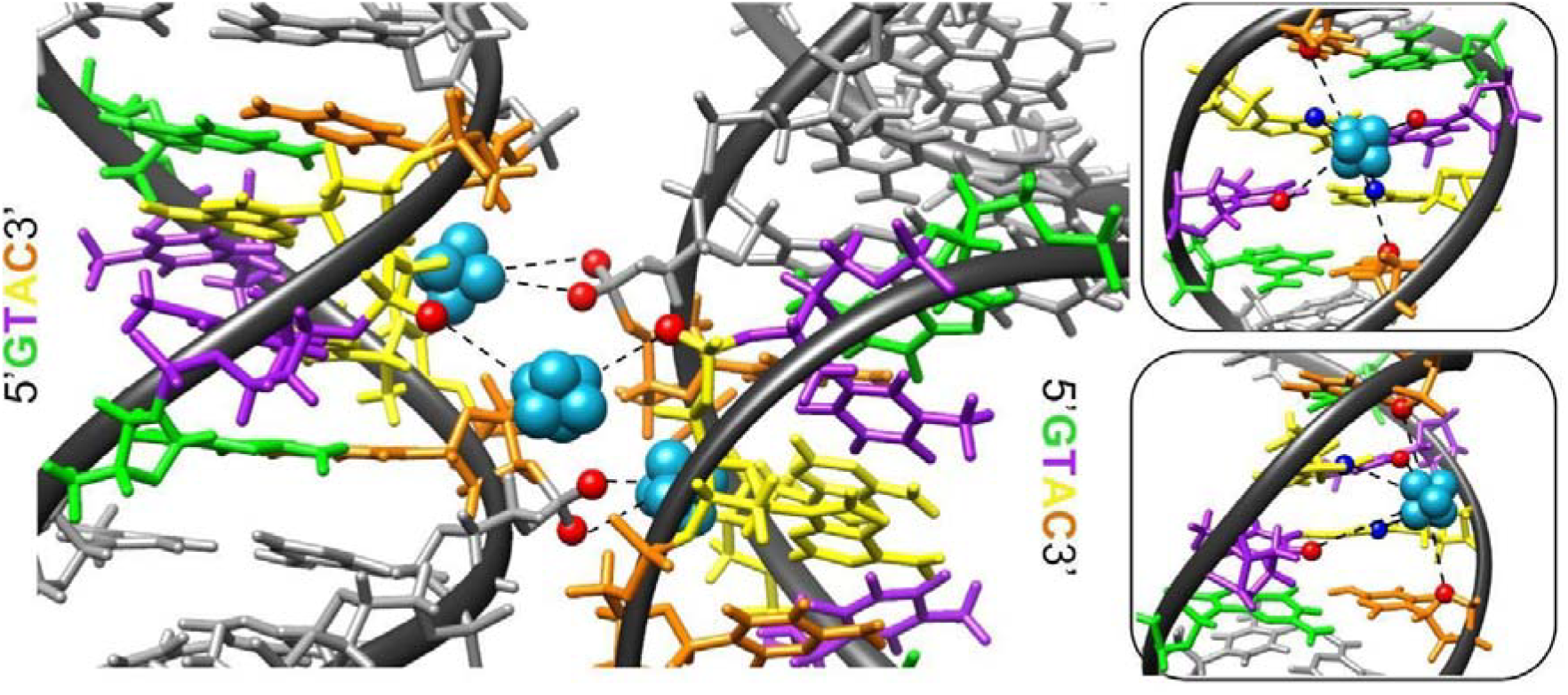
Molecular mechanism of Ni^2+^-mediated DNA-DNA condensation. Detailed views of DNA-ion-DNA bridges established in the 5’GTAC3’ motif from the pairing of the fragment 2 with itself (replica 2). The two ions located inside the minor groove (as shown in zooms) present the longest resident times, while the third one between backbones is more dynamic (Table S2). The sequence motif 5’-GTAC-3’ is highlighted in colour: A is represented in yellow, T in purple, C in orange, and G in green. Oxygen atoms are depicted in red and nitrogen atoms in blue. Zooms illustrate Ni^2+^ ions inside the minor groove from various perspectives. Dashed lines denote interactions between Ni^2+^ and DNA atoms. See Movie S2 of the simulation.

### A similar groove-alignment mechanism operates under physiological divalent-ion conditions

To determine whether the groove alignment mechanism is generalizable to physiological divalent ions, we performed simulations of DNA pairing in the presence of Mg^2+^ or Ca^2+^ ions. We initially carried out simulations at elevated ion concentrations (100 mM) to match the conditions used in Ni^2+^ simulations. Following the successful formation of parallel DNA pairings, we subsequently performed additional simulations under more physiological solvent conditions (150 mM KCl with 20 mM MgCl_2_ or CaCl_2_) (Table S1). These concentrations align closely with the physiological ranges of Mg^2+^ and Ca^2+^ reported for chromatin. In interphase nuclei, Ca^2+^ concentrations are typically 4–6 mM but increase to 12–24 mM in metaphase chromosomes, where meiotic homologous pairing occurs. Similarly, Mg^2+^ concentrations rise from 2–4 mM in interphase nuclei to 5–17 mM in metaphase chromosomes^37,52^. In practice, the DNA molecules inside the simulation solvation box were partially or fully neutralized by Mg^2+^ or Ca^2+^ ions, followed by the addition of ions to achieve a background concentration of 150 mM KCl (see Methods).

For DNA pairing in the presence of Mg^2+^ or Ca^2+^, we focused on fragments from the central region of the sequence, which exhibit a higher pairing probability in AFM experiments (Fig. 2). Although the proportion of pairing events was lower than that observed with nickel, pairing was nevertheless detected in two of four simulations at 100 mM Ca^2+^ and three of four simulations at 100 mM Mg^2+^ (Fig. 5, S10 and S11). Under physiological solvent conditions, we modelled the self-pairing of fragment 1 in the presence of Ca^2+^ and fragment 2 in the presence of Mg^2+^, guided by the pairing propensities observed in the 100 mM simulations (Fig. 5). Under these conditions, parallel pairing was observed in only two of six simulations with Ca^2+^ and one of six simulations with Mg^2+^ (Fig. 5, S10 and S11).

**Figure 5.**
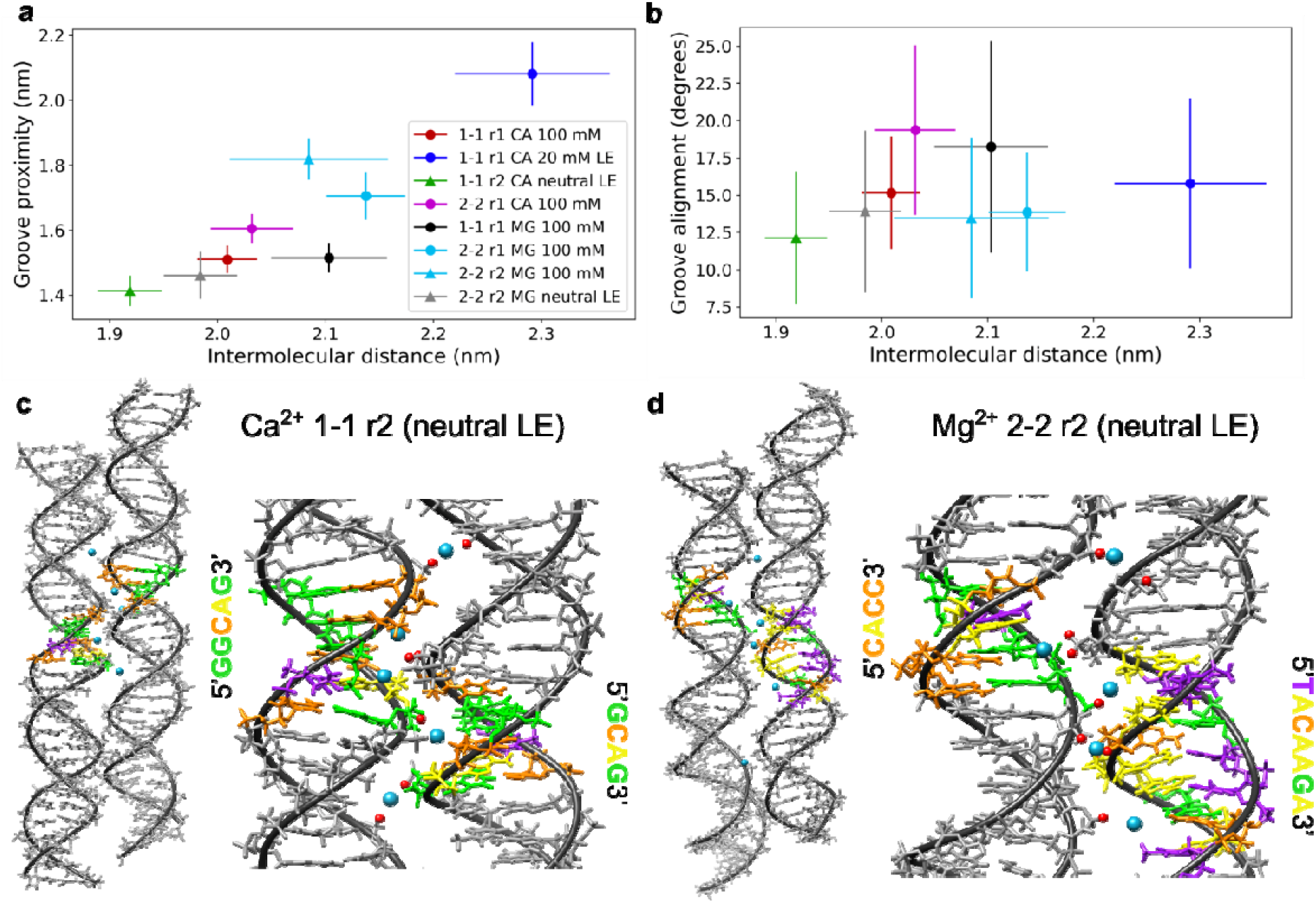
Molecular mechanism of DNA-DNA condensation mediated by Ca^2+^ and Mg^2+^. **a-b**. Strength of intermolecular DNA pairing as a function of the distance (**a**) and angle (**b**) between the minor grooves of adjacent duplexes. DNA pairings are coloured according to the interacting fragments (1 or 2), with replicas 1 and 2 (r1 and r2) distinguished by shape. Only simulations exhibiting parallel pairing are included, including: (i) 100 mM randomly distributed divalent cations (100 mM), (ii) 20 mM divalent cations placed at the lowest-energy favourable positions (20 mM LE), and (iii) charge-neutralised DNA systems with divalent cations placed at the lowest-energy favourable positions (neutral LE). No parallel pairing was observed in 20 mM randomly distributed divalent ions. Values represent averages over the final 100 ns of each simulation and error bars indicate standard deviations. **c–d**, Representative structures of the simulations showing the strongest DNA–DNA pairing in Ca^2+^ (1–1 r2 neutral LE) (**c**) and Mg^2+^ (2–2 r2 neutral LE) (**d**). Structures for the remaining Ca^2+^- and Mg^2+^-solvated simulations are shown in Fig. S10. Zooms highlight bridging ions (in cyan) with residence times >60%. Sequences hosting ions within the grooves are colour-coded by nucleotide identity: A, yellow; T, purple; C, orange; and G, green. Backbone oxygen atoms involved in ion bridging are shown in red. See Movies S5 and S6 of the simulations and Table S2 for more information regarding bridging ions.

We found that paired DNA duplexes could achieve intermolecular distances as low as those observed with nickel (<2.2 Å) and adopted a similar structure with highly aligned grooves (Fig. 5 and S10). In particular, we found a strong correlation of 0.90 between the distances of neighbouring minor grooves and intermolecular distances, mirroring the high correlation found in Ni^2+^ (0.81, see Fig. 3 and 5). Together, these findings suggest that the overall structure of DNA pairing mediated by Mg^2+^ and Ca^2+^ is similar to that of Ni^2+^ simulations, which in turn is consistent with the high-resolution AFM images (Fig. 1).

### Sequence motifs driving pairing depend on the divalent cation

The differences in each ion type’s ability to drive pairing stem from distinct ion atmospheres. Unlike Ni^2+^, Ca^2+^ and Mg^2+^ formed numerous bridging interactions between the two duplexes through a variety of binding configurations (Fig. 5, S12 and Movies S5–S8). Some ions were positioned within the minor groove of one duplex while simultaneously interacting with the backbone of the second duplex, resembling the bridging mode observed for Ni^2+^. In addition, other ions mediated pairing through direct backbone–backbone bridging interactions between the two DNA molecules (Fig. 5, S12, Table S2). Notably, Mg^2+^ ions localized within the major groove were also observed to mediate bridging interactions with the backbone of the adjacent duplex. While divalent ion binding inside the major groove was detected, consistent with previous crystallographic studies and simulations^25,51,53,54^, these interactions were only capable of driving ionic bridges in the case of Mg^2+^, failing to support effective inter-duplex bridging in the case of Ca^2+^ and Ni^2+^.

We observed that optimum pairing requires groove alignment driven by ionic interactions which can be sequence dependent. Simulations show that the sequence preference for ionic stabilization varies depending on the type of divalent ion. This could be due to the different propensities of each ion type to penetrate inside the grooves defined by distinct sequences. Overall, we noted that Ni^2+^ drives DNA pairing through a limited number of ions positioned inside the minor groove of the GTAC motif (Fig. 4, S9). Ca^2+^, on the other hand, allows for a larger number of bridging ions and presents a greater flexibility in sequence for locating ions in their minor groove, thus defining multiple pairing hotspots (Fig. 5, S12, Movies S5, S7). While Mg^2+^ presents a profile highly similar to Ca^2+^, its flexibility in defining pairing hotspots is further enhanced by its ability to coordinate ions within the major groove (Fig. 5, S12, Movies S6, S8). This additional coordination enables the formation of a continuous pairing “zipper,” where minor-groove bridging can be seamlessly extended by major-groove interactions. Overall, these molecular mechanisms explain the preference for pairing homologous sequences over non-homologous ones, as maintaining this orientation in long DNA molecules would necessitate the elimination of structural heterogeneity.

## Discussion

Here, we provide the first direct structural evidence for ion-mediated DNA–DNA pairing by integrating AFM with all-atom simulations. Although models such as the “electrostatic DNA zipper” or helical alignment mechanism have been proposed for more than two decades^27–29^, direct visualization of the underlying structural organization has remained experimentally elusive. By integrating AFM images with atomistic simulations, we show that paired DNA duplexes adopt a highly ordered groove-to-groove alignment stabilized by localized ionic bridges. The strong agreement between the groove alignment observed experimentally and the spontaneous self-assembly reproduced in solution by MD simulations demonstrates that this structural pairing is an intrinsic property of the DNA–ion atmosphere.

High-resolution AFM imaging revealed that duplex pairing is accompanied by clear alignment of major and minor grooves, consistent with the proposed DNA zipper model^27–29^. While homologous and non-homologous pairings were observed with similar frequency, contact analysis revealed discrete pairing hotspots concentrated primarily in the central region of the DNA molecules. These hotspots differed depending on the ionic environment, indicating that duplex association is not purely electrostatic but is instead modulated by sequence-dependent ion coordination. These observations are consistent with previous studies showing that divalent ion-mediated DNA attraction can occur between non-homologous molecules^26^ and that specific sequence motifs, such as those identified in *Neurospora crassa*, are capable of driving pairing^14^. Together, these findings suggest that sequence homology may not be required for the initial establishment of duplex–duplex contacts, but instead acts to stabilize and propagate pairing over longer distances by preserving structural registry between neighbouring helices.

Atomistic MD simulations reproduced the spontaneous condensation of DNA duplexes observed experimentally and revealed the molecular basis of these interactions. Across all divalent ions tested, the strongest pairing events occurred when neighbouring minor grooves became closely aligned, enabling ions to simultaneously coordinate both duplexes. Under AFM-like ionic conditions using Ni^2+^ as a counterion, the most stable interactions were associated with the GTAC motif, where ions localized within the minor groove formed long-lived bridges to the backbone of the adjacent duplex. These coordination sites generated highly stable intermolecular contacts with residence times extending over tens of nanoseconds, providing a structural mechanism through which local sequence motifs can nucleate DNA pairing.

Importantly, we show that this mechanism is not unique to Ni^2+^. Simulations performed with physiologically relevant divalent ions demonstrated that both Mg^2+^ and Ca^2+^ can also stabilize groove-aligned DNA pairings. Although the resulting structures closely resembled those observed with Ni^2+^, the ion atmospheres surrounding the duplexes differed substantially. Whereas Ni^2+^ pairing was dominated by a relatively small number of highly localized minor-groove ions, Mg^2+^ and Ca^2+^ produced a more diverse network of intermolecular bridges. These included minor-groove– backbone contacts, backbone–backbone interactions, and, in the case of Mg^2+^, additional bridging interactions involving ions localized within the major groove.

The coordination of Ca^2+^ and Mg^2+^ ions inside the minor groove has been previously observed in crystallographic structures, where bridging interactions between adjacent DNA duplexes were also detected as part of crystal packing arrangements^55,56^. Here, through atomistic simulations, we demonstrate that such side-by-side DNA–DNA configurations can also occur in solution. The long residence times observed in our simulations, together with their resolution in X-ray structures, indicate that these bridging divalent ions form stable intermolecular contacts capable of driving the DNA “zipper” mechanism through electrostatic interactions. Nevertheless, contributions from entropic effects and dynamic counterion fluctuations are also likely to play an important role^57^. It is well established that monovalent salts, which promote pairing in long DNA molecules, generate diffuse and dynamic counterion atmospheres around DNA^51,53,58^. We therefore propose a hierarchical mechanism in which stable divalent ion-mediated crosslinks nucleate local pairing events, while more dynamic ion interactions facilitate the propagation of pairing over longer length scales.

Together, our findings establish a structural framework for the DNA “zipper” model by demonstrating that sequence-dependent divalent ion coordination within DNA grooves underpins inter-duplex recognition. We propose a hierarchical model in which stable ion-mediated contacts nucleate local pairing, while sequence homology promotes the maintenance and propagation of pairing across extended regions of DNA. More broadly, these results provide a mechanistic basis for understanding how DNA duplexes can recognize and associate with one another in the absence of proteins, with potential implications for chromosome condensation, homologous chromosome pairing, and higher-order genome organization.

## Data availability

Data from this article is available at the York Research Database (DOI: 10.15124/803e24ad-328f-4df8-8885-4736192bfc85).

## Supplementary Data

Supplementary Data is available at NAR online.

## Acknowledgements

This work was supported by the CONACYT agency of the Mexican government (scholarship no. 291163) and by EPSRC grants EP/V027395/1 (V.V.), EP/N027639/1 and EP/Y008693/2 (A.N.), as well as the UKRI Future Leaders Fellowship MR/W00738X/1 (T.E.C. and A.L.B.P.). We gratefully acknowledge the Henry Royce Institute for Advanced Materials (EP/R00661X/1, EP/S019367/1, EP/P02470X/1, and EP/P025285/1) and thank Xinyue Chen for providing access to and support with the Dimension FastScan through Royce@Sheffield. We further acknowledge funding from EPSRC grants EP/T022205/1, EP/X035603/1, EP/P020259/1, EP/T022167/1 and University of York for computational resources.

## Author contributions

T.E.C, V.V A.L.B.P and A.N conceived and designed the experiments. T.E.C, D.E.R and A.L.B.P. conducted AFM experiments. T.E.C and D.E.R analysed AFM data. V.V conducted MD simulations and V.V and A.N. analysed them. T.E.C, V.V, A.L.B.P and A.N wrote the manuscript.

